# OmiCoreTumorDetector: an open, molecularly validated model for mapping tumour regions in colorectal cancer H&E sections

**DOI:** 10.64898/2026.09.21.753083

**Authors:** Niwase Shamim, Akihisha Fujiyama

**Affiliations:** OmiCore Inc., Fukuoka, Japan; Graduate School of Engineering, Kyushu University, Fukuoka, Japan

**Keywords:** computational pathology, colorectal cancer, H&E, tumour segmentation, stain normalisation, domain shift, calibration, spatial transcriptomics, Visium HD, open-source software

## Abstract

Defining tumour regions on haematoxylin and eosin (H&E) sections is a routine first step in spatial-omics studies, yet it is usually done by hand and is difficult to reproduce. We present OmiCoreTumorDetector, an openly licensed model that maps tumour-enriched regions in colorectal cancer (CRC) H&E sections and exports them as QuPath-compatible annotations. The released model (`omicore-tumordetector-crc-he-v0.1`) is an ensemble of three convolutional classifiers trained on 100,000 public tissue tiles, combined with Macenko stain normalisation at inference. During development we found that the main obstacle to reuse was calibration under stain-domain shift rather than discrimination: a single model kept an area under the ROC curve (AUROC) of 0.955 on unseen slides while its sensitivity at the conventional 0.5 threshold fell to 0.48. Training on non-normalised tiles raised tumour AUROC on an independently collected tile set from 0.836 to 0.992, and normalising at inference reduced false-positive tumour area in normal-adjacent tissue by 16-to 26-fold. On five 10x Visium HD CRC sections that share no material with the training data, the released model called 27.7–48.5% of tissue as tumour in three carcinoma sections and 0.08% and 1.82% in two normal-adjacent sections, exporting no tumour region from either normal section. On the carcinoma section with matched single-cell-resolution transcriptomics, agreement with transcriptome-derived tumour-cell identities reached an AUROC of 0.985 (95% spatial-block bootstrap CI 0.975– 0.993). The image model never observes gene expression, so this is orthogonal evidence. The model localises tumour-enriched regions at 112 µm resolution; it does not identify individual malignant cells and has not yet been validated across scanners, institutions or histological variants. Code, weights and evaluation are released under Apache-2.0 and installable with `pip install omicoretumordetector`.

## INTRODUCTION

Spatially resolved omics now measure gene and protein expression in intact tissue sections ^1–4^. Sequencing-based platforms have reached subcellular bin sizes ^5,6^, imaging-based platforms profile hundreds to thousands of transcripts in single cells ^7,8^, and multiplexed protein imaging resolves tens of proteins per cell ^9–12^. Studies of solid tumours on any of these platforms almost always begin with a morphological question: which part of the section is tumour? The answer determines which regions are profiled, how tumour core, tumour edge and adjacent tissue are compared^13–15^, and how results are interpreted. In practice the region is drawn by hand. This is slow, depends on the operator and is difficult to report reproducibly.

Deep learning on routine H&E histology is now well established^16–18^, and general-purpose pathology foundation models are emerging ^19,20^. Tile-level tissue classifiers for colorectal H&E have been available since the release of the NCT-CRC-HE-100K dataset^21,22^, and they reach very high accu-racy on held-out tiles from the same source. That accuracy has not translated into tools that groups routinely apply to their own slides. Differences in staining protocol and scanner between laboratories are a well-documented source of performance loss in computational pathology^23–25^, and colour normalisation ^26,27^ and colour augmentation^25^ are the usual countermeasures. Less attention has been paid to how such shifts affect the *calibration* of predicted probabilities ^28^. In a region-mapping tool this matters directly, because the output is produced by thresholding those probabilities.

Here we treat robustness to stain-domain shift, rather than headline tile accuracy, as the design target. We build an ensemble tumour-region mapper for colorectal H&E and test it on whole sections from a separate study. We validate it against a modality the model cannot see: transcriptome-derived cell identities from the same tissue section^6^. We release the model as an installable Python package with QuPath^29^ export, and we report the approaches that did not work alongside those that did.

## RESULTS

### Model construction and held-out benchmarks

We trained three nine-class tissue classifiers (adipose, background, debris, lymphocytes, mucus, smooth muscle, normal mucosa, cancer-associated stroma and adenocarcinoma epithelium) on the 100,000-tile NCT-CRC-HE-100K dataset^22^: one ConvNeXt-Tiny^30^ on the Macenko-normalised release and one ConvNeXt-Tiny and one EfficientNetV2-S^31^ on the non-normalised (NONORM) release of the same tiles (Fig. 1a,b; Methods). The released model averages the three classifiers’ probabilities after Macenko normalisation of each input tile. The tumour-epithelium (TUM) probability is used as the tumour score.

**Figure 1.**
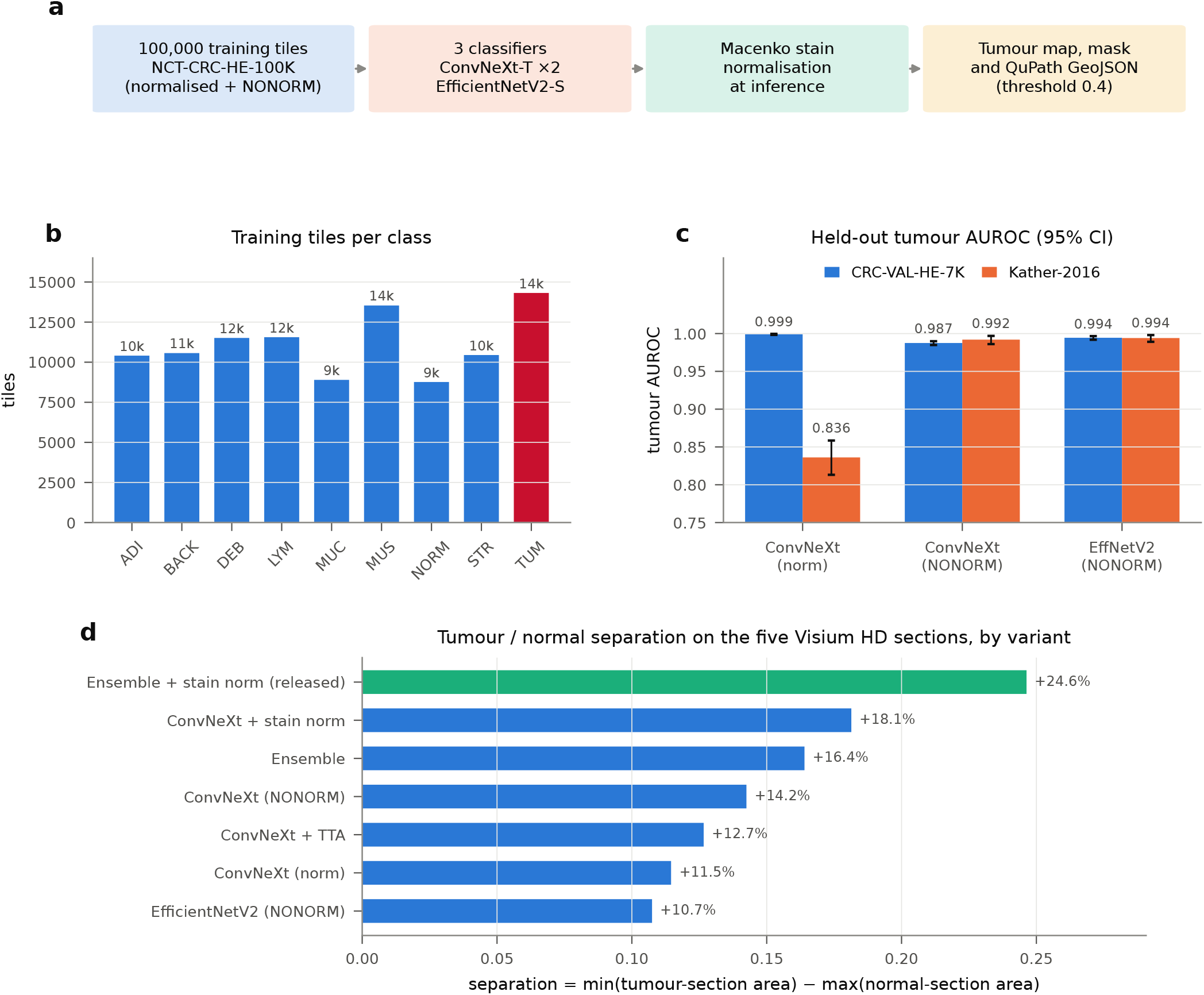
Construction and benchmarking of OmiCoreTumorDetector. **a**, Overview of the pipeline. Three nine-class tile classifiers were trained on NCT-CRC-HE-100K (100,000 tiles of 224 × 224 px at 0.5 µm per pixel): a ConvNeXt-Tiny on the Macenko-normalised release, and a ConvNeXt-Tiny and an EfficientNetV2-S on the non-normalised (NONORM) release of the same tiles. At inference, every tissue tile of a section is Macenko-normalised, scored by all three classifiers, and their softmax outputs are averaged; the tumour (TUM) probability is thresholded at 0.4 to give a tumour mask and QuPath-readable GeoJSON polygons (Methods, Algorithm 1). **b**, Number of training tiles in each of the nine tissue classes (total 100,000). ADI, adipose tissue; BACK, background; DEB, debris; LYM, lymphocytes; MUC, mucus; MUS, smooth muscle; NORM, normal colon mucosa; STR, cancer-associated stroma; TUM, colorectal adenocarcinoma epithelium (red). **c**, Tumour-versus-rest AUROC of each ensemble member on two held-out tile sets: CRC-VAL-HE-7K (blue; 7,180 tiles from 50 patients absent from training, same tissue bank) and Kather-2016 (orange; 4,375 tiles from a separately collected and digitised collection). Bars show point estimates (values above bars) and whiskers 95% bootstrap confidence intervals (1,000 tile resamples). The ConvNeXt trained on normalised tiles is near-perfect on CRC-VAL-HE-7K (0.999) but falls to 0.836 on Kather-2016, whereas both models trained on NONORM tiles stay at or above 0.992. **d**, Separation between the three carcinoma and two normal-adjacent Visium HD sections for each inference variant, defined as the smallest carcinoma-section tumour-area fraction minus the largest normal-adjacent tumour-area fraction at threshold 0.5 (Eq. 14); values above zero mean the two groups do not overlap. The released configuration (ensemble with stain normalisation, green) gives the largest separation, +24.6 percentage points. TTA, test-time augmentation over four flips. Values for every variant are listed in Supplementary Table S2.

We evaluated each classifier on two held-out tile sets (Fig. 1c; Supplementary Table S1). CRC-VAL-HE-7K contains 7,180 tiles from 50 patients who do not appear in the training set. It comes from the same tissue bank and was prepared in the same way, so it tests generalisation to new patients rather than to a new laboratory. On it, the ConvNeXt trained on normalised tiles reached a tumour AUROC of 0.999 (95% CI 0.998–1.000). The Kather-2016 texture collection^32,33^ was collected and digitised separately (0.495 µm per pixel, 150 px tiles). On this set the same model fell to 0.836 (0.813– 0.859), whereas the identically configured ConvNeXt trained on NONORM tiles reached 0.992 (0.986–0.997) and the EfficientNetV2 0.994 (0.989–0.998). Internal validation accuracy on a random 15% split of the training data was 99.8%. We do not use this figure: NCT-CRC-HE-100K carries no patient identifiers (the code in each file name is unique to each tile), so the split very probably places tiles from the same patient on both sides.

### Stain-domain shift causes a calibration failure rather than a ranking failure

The target sections were measurably outside the training colour domain. Tissue tiles from the Visium HD sections were lighter and pinker than NCT tiles, at a mean RGB distance of 56.3 from the training colour mean. Macenko normalisation reduced this distance by 51%, to 27.6 (Fig. 2a,b).

**Figure 2.**
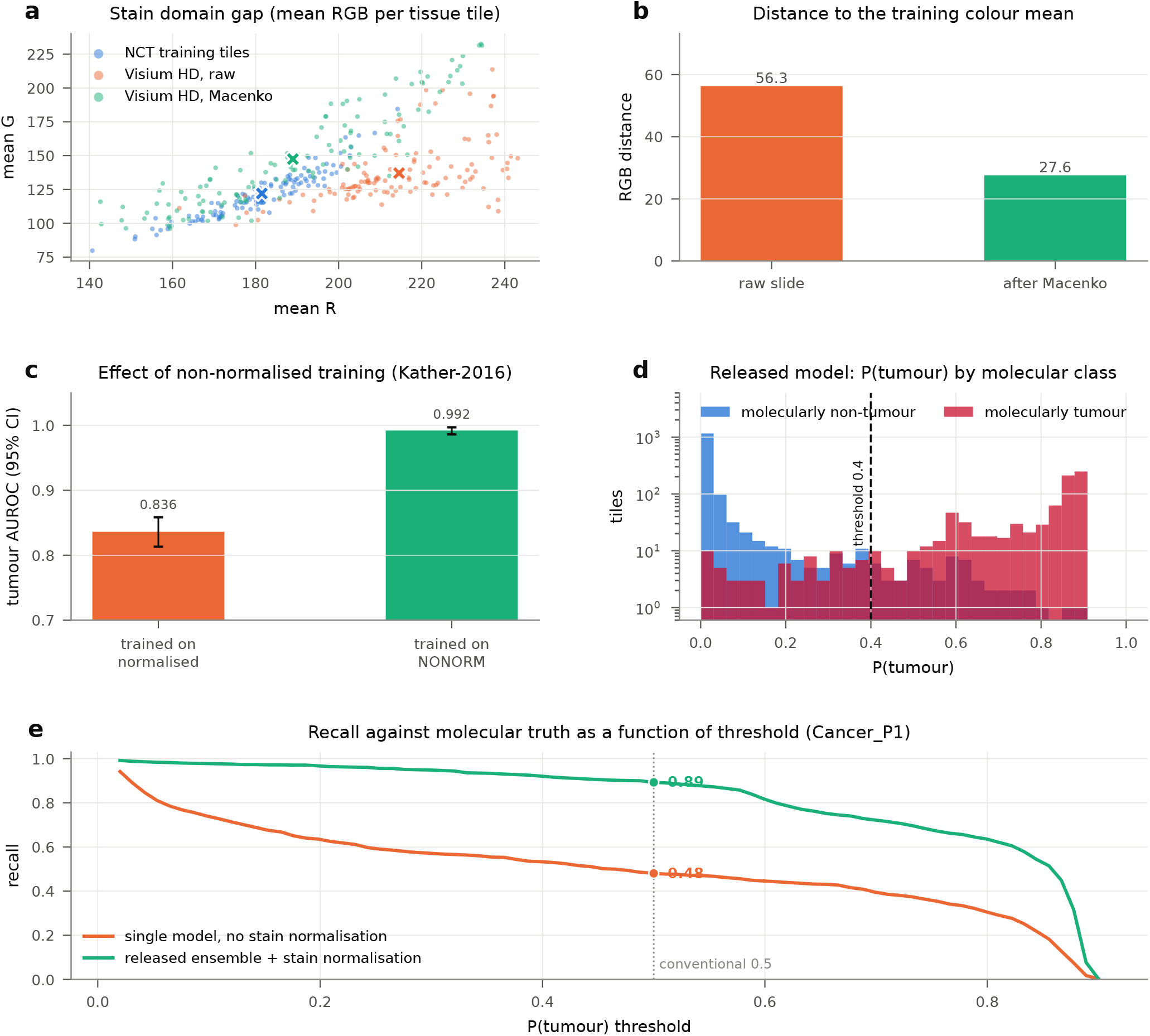
Stain-domain shift is a calibration failure, and how the released model corrects it. **a**, Mean red versus mean green intensity of individual tissue tiles: 120 NCT-CRC-HE-100K tumour tiles (blue; the training domain), 120 tissue tiles sampled at random from the raw Cancer_P1 Visium HD H&E image (orange), and the same 120 tiles after Macenko normalisation (green). Crosses mark the mean of each group. The raw Visium HD tiles are lighter and pinker than the training tiles; normalisation moves them towards the training domain. **b**, Euclidean distance, in mean-RGB space, between the Visium HD tile mean and the NCT tile mean before (56.3) and after (27.6) normalisation, a 51% reduction. **c**, Tumour-versus-rest AUROC on the Kather-2016 set for two identically configured ConvNeXt-Tiny models that differ only in their training tiles (Macenko-normalised or non-normalised); whiskers, 95% bootstrap confidence intervals (1,000 resamples). **d**, Distribution of the released model’s tumour probability for Cancer_P1 tiles containing at least 20 transcriptomically labelled cells, split by molecular class: tumour-dominant tiles (≥50% of cells assigned a tumour identity; *n* = 859, red) and all other tiles (*n* = 1,448, blue). The *y*-axis is logarithmic; the dashed line marks the released threshold of 0.4. **e**, Sensitivity against the transcriptomic reference as the probability threshold is varied, for a single ConvNeXt trained on normalised tiles and applied to raw slides (orange) and for the released ensemble with stain normalisation (green). Dots mark the conventional threshold of 0.5, where sensitivity is 0.48 and 0.89, respectively. Because the single model’s AUROC is 0.955, its low sensitivity at 0.5 reflects compressed (miscalibrated) probabilities rather than poor ranking of tiles.

The consequence was a failure of calibration, not of discrimination. Applied to raw target slides, the single classifier trained on normalised tiles ranked tiles almost as well as the released model against the transcriptomic reference (AUROC 0.955; Supplementary Table S2). However, its predicted probabilities were compressed towards zero, and at the conventional threshold of 0.5 it recovered only 48% of molecularly tumour-dominant tiles (Fig. 2e). Such a model looks highly precise (precision 0.981) while missing half of the tumour, and nothing in its output signals the problem.

The two design choices address different halves of this problem (Supplementary Table S2). Training on NONORM tiles improved cross-collection discrimination (Fig. 2c) and sensitivity. Normalising at inference restored calibration and specificity: for the single ConvNeXt at threshold 0.5 it reduced the tumour area called in normal-adjacent tissue from 1.06% to 0.04% (Normal_P3) and from 4.78% to 0.30% (Normal_P5). Combining the two in the released ensemble gave the best result on both independent criteria we used: section-level separation (Fig. 1d) and agreement with transcriptomic identities (described below). At 0.5 its sensitivity against the transcriptomic reference was 0.893, compared with 0.481 for the single model on raw slides (Fig. 2e).

### Tumour-region mapping on carcinoma and normal-adjacent sections

We applied the released model to five 10x Visium HD formalin-fixed, paraffin-embedded sections from the study of Oliveira et al. ^6,34^: three primary CRC sections (patients P1, P2 and P5) and two normal-adjacent sections (P3 and P5) (Fig. 3; Table 1). The model called 32.3%, 48.5% and 27.7% of tissue as tumour in the three carcinoma sections, and 0.08% and 1.82% in the two normal-adjacent sections. After the minimum-area filter used for annotation export (Methods), neither normal-adjacent section produced a single tumour region, and the smallest carcinoma section fraction exceeded the largest normal-adjacent fraction by 25.8 percentage points.

**Table 1.**
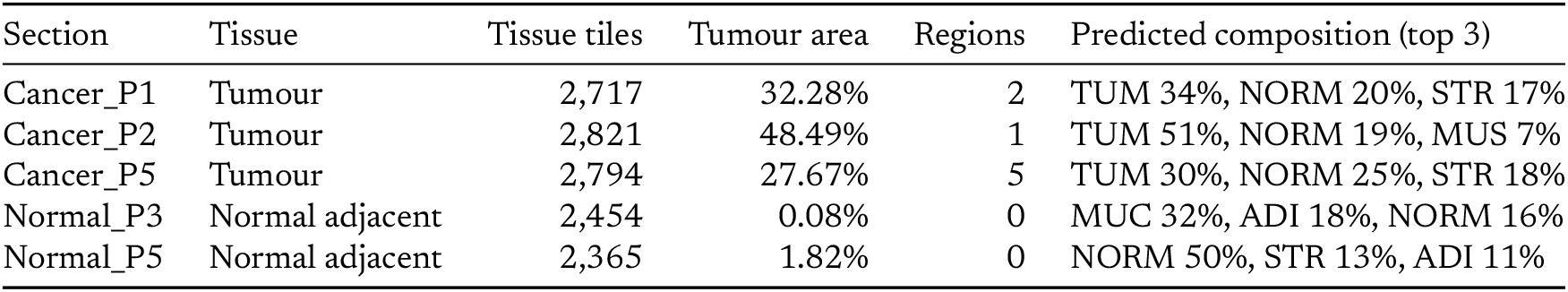
Released model on the five Visium HD sections. (threshold 0.4). Results from the package installed from the public index; identical to the development pipeline. Regions: tumour polygons exported after smoothing and the minimum-area filter. Composition: the three most frequent predicted tissue classes.

**Figure 3.**
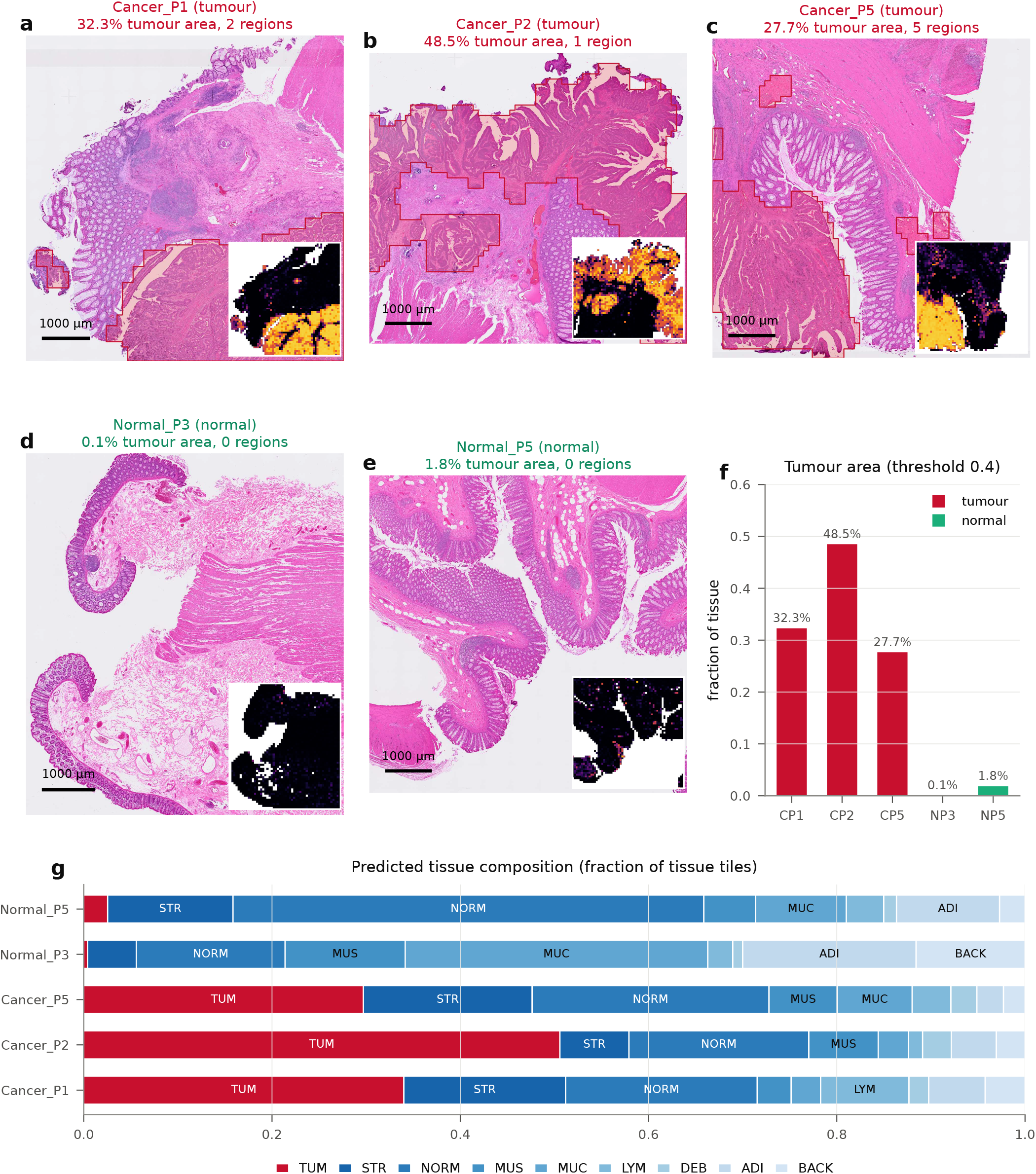
Tumour-region mapping on three carcinoma and two normal-adjacent sections. **a–e**, H&E images of the five 10x Visium HD FFPE sections of Oliveira et al. ^6^, shown at 2 µm per pixel (downsampled four-fold from the 0.5 µm per pixel analysis resolution). The red outline and light red tint mark the tumour regions exported by the released model: tiles with ensemble tumour probability ≥0.4, smoothed by a 3 × 3-tile morphological closing and opening, with regions smaller than 50,000 px^2^ (12,500 µm^2^) removed (Eqs. 10–11). Panel titles give the fraction of tissue tiles called tumour and the number of exported regions. Insets (lower right) show the per-tile tumour probability (black, 0; yellow, 1; white, glass, not scored). **a–c**, Primary colorectal adenocarcinomas from patients P1, P2 and P5. In **b**, the region contains a hole where an enclosed island of normal mucosa was excluded. **d, e**, Normal-adjacent tissue from patients P3 and P5, in which no region was exported. Scale bars, 1 mm. Tissue images are embedded at 1,391–1,586 ppi at printed size. **f**, Fraction of tissue tiles with tumour probability ≥0.4 in each section (C, carcinoma; N, normal-adjacent). **g**, Predicted tissue composition: the fraction of tissue tiles whose most probable class is each of the nine tissue classes, with TUM in red and the other classes in shades of blue (abbreviations as in Fig. 1b).

The predicted tissue composition was consistent with the histology (Fig. 3g). Tumour epithelium, stroma and normal mucosa dominated the carcinoma sections, whereas normal mucosa, mucus, adipose tissue and stroma dominated the normal-adjacent sections. In Cancer_P2 the exported region contains a hole: an island of 19 tissue tiles (about 0.24 mm^2^), 14 of them classified as normal mucosa, fully enclosed by tumour tiles and left out of the tumour region rather than absorbed into it (Fig. 3b).

A negative control is necessary but not sufficient: a model that never predicted tumour would also pass it. The evidence rests on both sides holding at once, and on the molecular comparison below.

### Agreement with transcriptome-derived tumour-cell identities

The Cancer_P1 section carries single-cell-resolution transcriptomics from the same tissue. In a companion analysis of these data, nuclei were segmented from the H&E image, 2 µm bins were assigned to cells with bin2cell ^35^, and each cell received an identity from a classifier trained on the study’s matched single-cell reference (Methods). We projected 202,182 cells onto the model’s tile grid. Of the 2,307 tiles containing at least 20 cells, 859 were molecularly tumour-dominant (at least half of their cells assigned a tumour identity).

The two independent maps localised the same compartment (Fig. 4a,b). Tile-level agreement gave an AUROC of 0.985 (95% spatial-block bootstrap CI 0.975–0.993) and a Spearman correlation of 0.769 between the H&E tumour probability and the molecular tumour-cell fraction (Fig. 4c,d; Table 2). At the released threshold of 0.4, precision was 0.935 (0.909– 0.959), sensitivity 0.921 (0.845–0.966) and specificity 0.962 (0.941–0.979). Because neighbouring tiles are not independent, we resampled contiguous 8 × 8-tile blocks rather than individual tiles. This widens the sensitivity interval to more than twice the tile-level bootstrap width (Methods). The AUROC was stable across definitions of a molecularly positive tile, ranging from 0.977 to 0.990 (Supplementary Table S3).

**Table 2.**
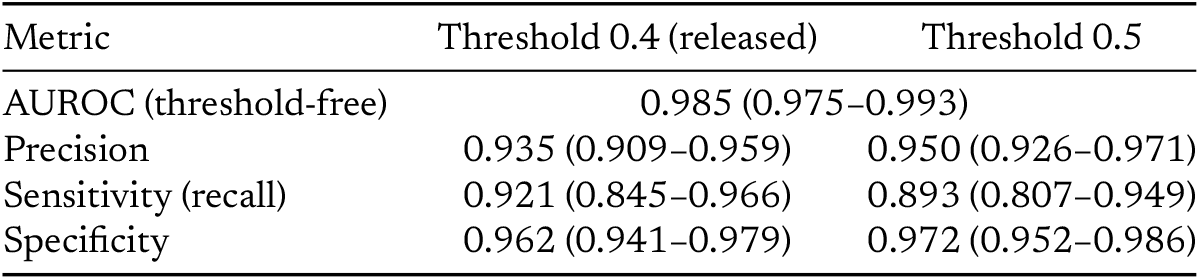
Agreement with transcriptome-derived tumour-cell identities. (Cancer_P1; 2,307 tiles with ≥20 cells, 859 molecularly tumour-dominant). Values in parentheses are 95% spatial-block bootstrap intervals. The 0.4 threshold was selected on this section, so threshold-dependent values in that column are optimistic; the AUROC is threshold-free.

**Figure 4.**
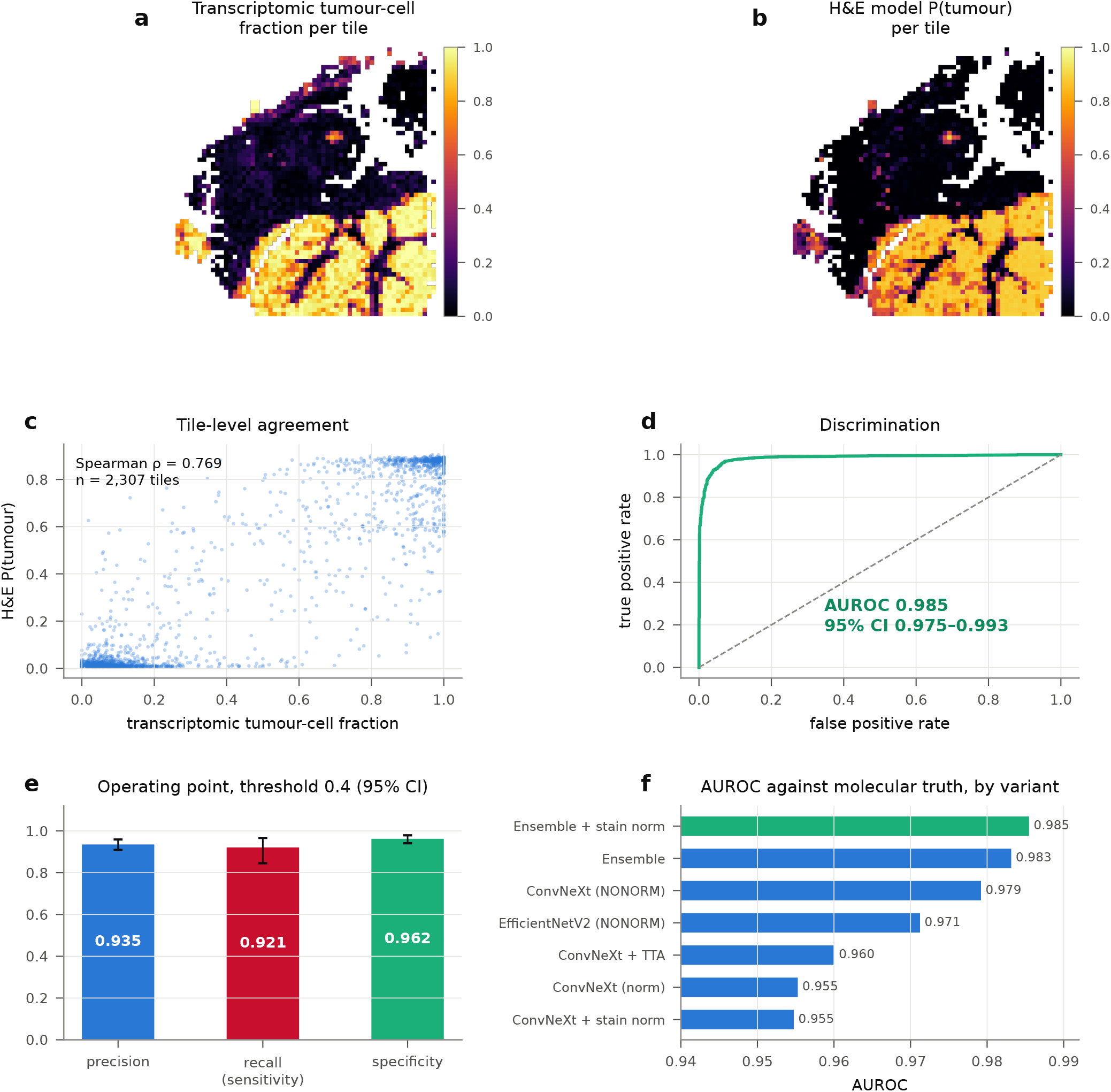
Validation against transcriptome-derived cell identities. All panels show Cancer_P1, the only section with matched single-cell-resolution transcriptomics. **a**, Fraction of cells in each 224-px (112 µm) tile that were assigned a tumour identity by a logistic-regression classifier trained on the study’s Chromium Flex single-cell reference (Methods); 202,182 cells were projected onto the tile grid, and tiles with fewer than 20 cells are left blank. **b**, Tumour probability from the released H&E model for the same tiles; the image model never observes gene expression. **c**, Tile-level relationship between H&E tumour probability and transcriptomic tumour-cell fraction (*n* = 2,307 tiles with ≥20 cells; Spearman *ρ* = 0.769). **d**, ROC curve of the H&E tumour probability for separating molecularly tumour-dominant tiles (≥50% tumour cells, *n* = 859) from all others (*n* = 1,448). AUROC 0.985 with a 95% confidence interval of 0.975–0.993 from a spatial block bootstrap (57 blocks of 8 × 8 tiles, 2,000 resamples; Methods); the dashed diagonal marks chance. **e**, Precision, sensitivity and specificity at the released threshold of 0.4, with 95% spatial-block bootstrap confidence intervals. Because the threshold was chosen on this section, these values are optimistic; the AUROC in **d** does not depend on a threshold. **f**, AUROC against the transcriptomic reference for every inference variant (Supplementary Table S2), with the released configuration in green.

Two caveats bound this result. First, the reference is itself a model output. The transcriptomic classifier assigned the correct broad lineage to 87.8% of cells from a held-out patient in the single-cell reference, so the reference carries label noise that caps the agreement any image model could reach. Second, the 0.4 operating point was chosen on this same section, so the threshold-dependent metrics are optimistic. The AUROC is threshold-free and unaffected.

### Histology at the boundary and the hardest negatives

Full-resolution crops at the 0.5 µm per pixel scale the model operates on showed the expected morphology on each side of the predicted boundary (Fig. 5a–h). Crops inside the predicted regions showed irregular, crowded glands. Crops outside them showed regularly spaced crypts, and boundary crops showed the transition between the two.

**Figure 5.**
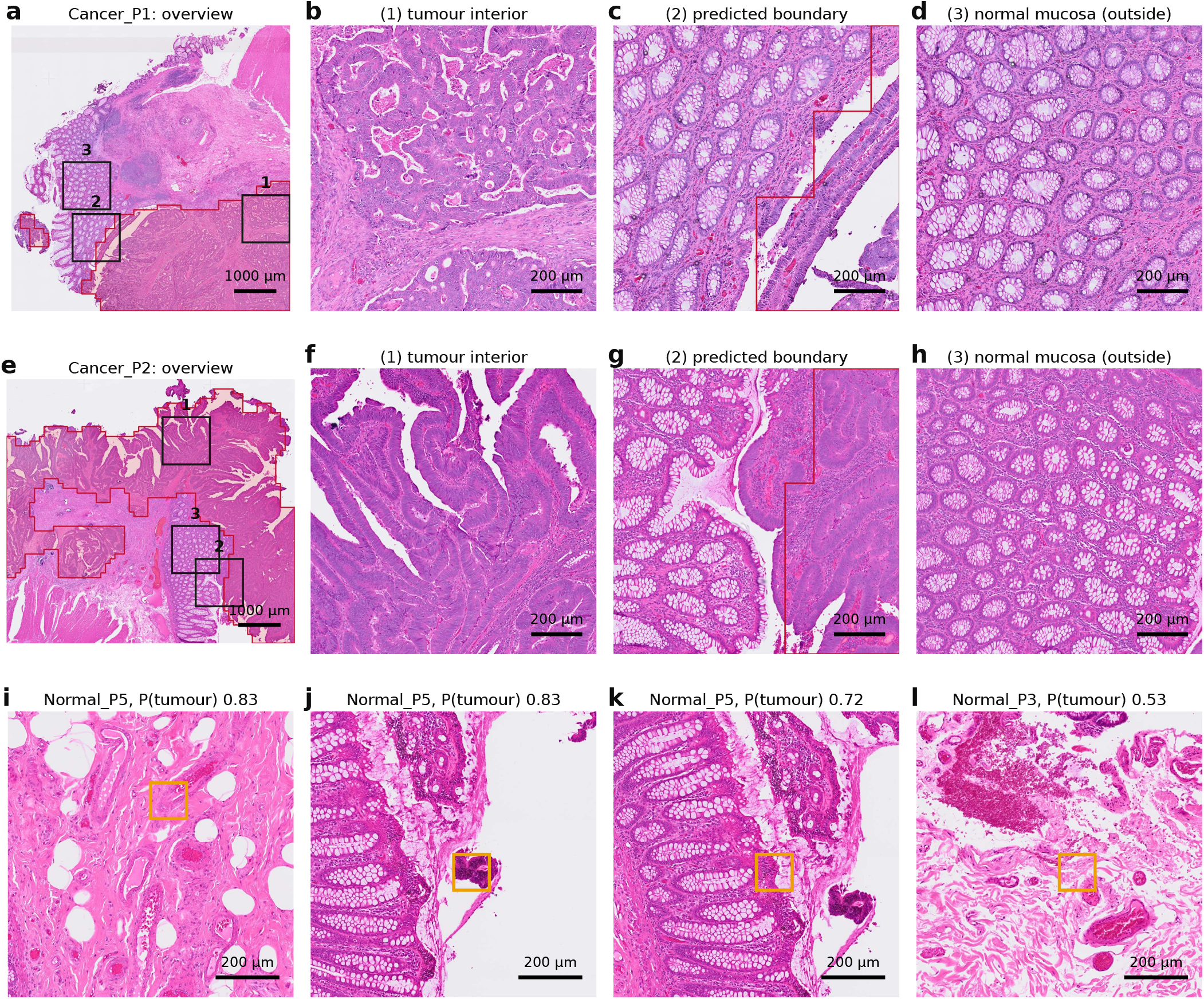
Histology at the model’s native resolution. **a, e**, Overviews of Cancer_P1 and Cancer_P2 (downsampled eight-fold) with the exported tumour region (red outline, light tint) and the positions of three crops (numbered boxes). Crop positions were chosen automatically from the tile maps, not by eye. **b, f**, Crop (1), the 10 × 10-tile window with the highest tumour coverage, inside the predicted region; it shows crowded, irregular glandular architecture. **c, g**, Crop (2), the window where the tumour mask meets tiles classified as normal mucosa; the red line is the exported boundary. **d, h**, Crop (3), the window with the highest fraction of tiles classified as normal mucosa and no tumour tiles, outside the predicted region; it shows regularly spaced crypts. Crops **b–d** and **f–h** are 2,240 × 2,240 px (1.12 mm across) at 0.5 µm per pixel, the resolution the model operates on, and are embedded at 1,350 ppi. **i–l**, The four highest-scoring tiles in the two normal-adjacent sections (yellow boxes, 224 px = 112 µm), each shown within 1,792 × 1,792 px (0.9 mm) of surrounding tissue; titles give the section and the tumour probability. None of these tiles formed an exported region. Scale bars, 1 mm (**a, e**) and 200 µm (all other panels).

We also examined the tiles in the normal-adjacent sections that received the highest tumour probability (Fig. 5i–l). The highest-scoring tiles (probability 0.83) lay at a tissue edge adjacent to a detached tissue fragment and within loose stromal and adipose tissue rather than within intact mucosa. None exceeded the minimum-area filter used for export. We present these as candidate failure modes, namely tissue edges, detached fragments and loose stroma, pending formal pathologist review.

### Software

The model is distributed as the Python package `omicoretumordetector` (Apache-2.0). Four lines of code map a section and write QuPath-readable GeoJSON:

~~~
from omicoretumordetector import TumorDetector
det = TumorDetector.from_pretrained()
res = det.predict(“section.tiff”, mpp=0.5)
det.to_geojson(res, “tumour.geojson”, threshold=0.4)
~~~

Input magnification is specified explicitly (`mpp`) and images are rescaled to the 0.5 µm per pixel training resolution, because silently mismatched magnification is a common cause of failure. Weights are downloaded on first use and verified against SHA-256 checksums. A 13,000 × 13,000 px section is processed in about 50 s on a single laptop GPU, including stain normalisation. We confirmed that the package installed from the public index reproduces the development pipeline exactly: tumour area agreed to within 0.004 percentage points and exported region counts were identical on all five sections.

## DISCUSSION

The central finding of this work is not that a convolutional ensemble can classify colorectal tissue, which has been established for several years^16,21,32^. It is that the barrier to reusing such models on another laboratory’s slides is calibration rather than capability. A model whose AUROC is essentially unchanged can lose half its sensitivity purely because the probability scale has shifted, and it does so quietly, presenting as high precision. Any group applying a tile classifier to its own material should therefore check the operating point before trusting thresholded output, even when the model’s discrimination has been validated elsewhere.

Two approaches that appeared promising were not adopted. Spatial smoothing of the probability map, motivated by the contiguity of tumour, changed the AUROC by at most 0.002 and reduced sensitivity in every configuration we tested (evaluated on an earlier single-model checkpoint). Label-free automatic thresholding (the triangle method) recovered sensitivity on tumour-bearing slides but fabricated tumour on tumour-free tissue: with the threshold fitted to each slide, the single ConvNeXt model called 12.6% of normal-adjacent tissue and 39.4% of an out-of-organ lung section colorectal tumour. The package retains it only behind an explicit opt-in, for delineating a tumour already known to be present. We report both, with their numbers, so that others need not repeat them.

### Uses in spatial transcriptomics

The model’s output is a reproducible, versioned tumour region per section, defined by fixed rules rather than by an annotator. We see five uses in spatial transcriptomics studies, summarised with their requirements and limits in Table 3.

**Table 3.** Proposed uses of OmiCoreTumorDetector in spatial omics studies. Each use relies only on the exported tumour polygons, the per-tile tumour probability or the per-tile class probabilities. ST, spatial transcriptomics; SP, spatial proteomics; ROI, region of interest.

| Use | Assay | What the model provides | Requirement or limit |
| --- | --- | --- | --- |
| Compartment labels | ST, SP | Tumour or non-tumour label for each spot, bin or cell inside or outside the polygons | Same coordinate frame as the H&E image; labels are region-level, not cell-level |
| Distance to boundary | ST, SP | Signed distance from each spot, bin or cell to the tumour boundary, binned into core, edge and distant tissue | Bands at least 112 $\mu\text{m}$ (one tile) wide |
| Orthogonal tumour check | ST | Image-based tumour region, independent of the expression reference, to compare with deconvolution or copy-number calls | Label noise in the expression-based reference limits agreement |
| Composition covariate | ST, SP | Fraction of tumour-called tiles per region or neighbourhood | The tile fraction approximates, but does not equal, tumour-cell purity |
| Capture-area and field selection | ST, SP | Tumour fraction and boundary location on a serial H&E section before the assay | Serial-section offset; H&E and assay sections must come from the same block |
| ROI selection for collection | SP | Rule-based ROIs (tumour, boundary band, distant tissue), exported to QuPath | ROIs follow the 112 $\mu\text{m}$ tile grid; review before collection |

First, *compartment labelling*. Every spot, bin or segmented cell whose coordinates fall inside the exported polygons can be labelled as tumour region, and the remainder as non-tumour tissue. The label is then available for compartment-level comparisons such as pseudobulk differential expression between tumour and adjacent tissue, applied identically to every section of a cohort. Because the GeoJSON polygons are in the pixel coordinates of the input H&E image, this needs only the image-to-array transform that Visium, Visium HD and in situ platforms already provide.

Second, *distance to the tumour boundary*. The signed distance from each spot, bin or cell to the nearest polygon edge turns the binary region into a continuous axis from tumour core through the boundary to distant tissue. Transcriptional programmes that differ between tumour core and leading edge^15^, at the tumour–microenvironment interface^14^ and in tumour-specific keratinocyte niches at the margin^13^ have been described along such axes. In colorectal cancer, immune hubs^36^, cellular neighbourhoods at the invasive front^37^ and the density of immune cells in the tumour core and invasive margin ^38,39^ all depend on position relative to the tumour. Because our boundary is resolved at 112 µm, distances should be binned in bands at least one tile wide. Distances measured on a finer scale than this are not supported by the model.

Third, *an orthogonal check on expression-based tumour calls*. Tumour identity in spatial data is commonly inferred from expression, by deconvolution against a single-cell reference ^40,41^ or by inference of copy-number alterations^42^. These calls depend on the reference and on the chosen parameters. An image-derived region does not share either dependency, so agreement between the two supports both, and disagreement localises tissue where one of them may be wrong. The comparison on Cancer_P1 reported here is one such check, and the per-tile agreement analysis can be repeated on any section with cell-level annotations.

Fourth, *a covariate for tissue composition*. Differences in expression between samples or regions are confounded by the proportion of tumour cells they contain, as shown for bulk tumour profiles ^43^. The fraction of tumour-called tiles in a region, or in the neighbourhood of a spot, can be included as a covariate so that changes in composition are not mistaken for changes within cells. Spatial clustering and resolution-enhancement methods that already use histology^44–46^ and models that predict expression from H&E ^47^ could use the mask in the same way, as a region-level prior or label.

Fifth, *experimental design*. Capture areas are small and costly. Running the model on a serial H&E section from each candidate block before the assay shows whether the planned capture area contains tumour, adjacent non-tumour tissue and the boundary between them, and allows the tumour fraction to be balanced across the samples of a study. A run takes about a minute per section on a laptop GPU.

### Uses in spatial proteomics

The same outputs apply to spatial proteomics. They matter most where regions of interest must be chosen before the measurement. In region-based digital spatial profiling ^48^, and in laser microdissection coupled to mass-spectrometry proteomics ^49^, the analyst chooses which regions to collect. The model can propose them by fixed rules, for example the tumour region, a band either side of its boundary, and tumour-free tissue at a stated distance. This reduces selection bias between operators and lets the rule be reported with the data. The polygons can be imported into QuPath ^29^ for review before collection.

Highly multiplexed protein imaging, by imaging mass cytometry ^9^, multiplexed ion beam imaging ^10,50^, CODEX ^11^ or cyclic immunofluorescence ^12,51^, often covers selected fields or tissue microarray cores rather than whole sections. The model can choose those fields from a whole-section H&E image and record where each field lies relative to the tumour. After acquisition, cells segmented from the protein image ^52,53^ can be assigned to the tumour region or to a boundary band. Neighbourhood analyses^37,50,54^ can then be stratified by compartment. In colorectal cancer, neighbourhoods at the invasive front have been linked to outcome^37^.

These uses depend on registering the H&E image to the protein image. When H&E staining is done on the same section after the protein assay, registration can be close to exact. When it is done on a serial section, the error is typically small at tissue scale but not at single-cell scale. Because the model resolves regions at 112 µm, it is suited to tissue-scale registration, but region edges should not be used to classify individual cells.

### Recommended practice

From these results, we suggest five practices for users. Record the pixel size of each image and pass it to the model. Check the operating point on a small subset of sections against a pathologist or a molecular reference before relying on thresholded output. Treat the boundary as a band of at least one tile rather than a line. Report the model version, threshold and minimum-area setting with the results. Do not use region membership as evidence that an individual cell is malignant; that requires cell-level evidence such as expression programmes or copy-number alterations ^42^.

### Limitations

The evaluation is small: three carcinoma and two normal-adjacent sections from a single study, of which one carcinoma section had matched transcriptomics. This demonstrates feasibility, not generalisation across patients, institutions or scanners. The transcriptomic reference is itself a classifier output with measurable label noise, and the operating threshold was selected on the section used to report threshold-dependent metrics. The model marks regions, not cells. Boundaries are resolved at the 224 px (112 µm) tile scale, so a called region necessarily contains stromal, immune and vascular cells alongside malignant glands, and isolated infiltrating glands at a margin may be missed. Tumour buds, single cells or clusters of up to four cells at the invasive front ^55^, are far below this scale and will not be detected. The evidence supports the claim that the model localises tumour-enriched regions; it does not support the claim that it identifies malignant cells. The model is colorectal-specific. Lung and prostate adenocarcinoma sections, used as out-of-organ negative controls, produced 8.9% and 1.7% spurious tumour area at the released threshold. Adenoma, dysplasia, inflammation, ulceration, necrosis, mucinous and signet-ring variants, and treatment-altered tissue were not represented in evaluation. The NCT normal-mucosa class also includes non-tumorous gastric tissue ^22^. The software is intended for research use only and is not a medical device.

### Future work

In order of value, the next steps are: independent pathologist annotation of invasive carcinoma on the same sections, scored with overlap and boundary-distance metrics rather than tile metrics; extension of the transcriptomic comparison to the remaining sections, which requires only re-running the existing annotation pipeline; external validation on unseen patients from several institutions and scanners, deliberately enriched for difficult non-neoplastic tissue; and replacement of the ImageNet-initialised backbones with pathology foundation models ^19,20^, which may improve robustness to staining and scanner differences.

## METHODS

### Training data

NCT-CRC-HE-100K comprises 100,000 non-overlapping 224 × 224 px tiles at 0.5 µm per pixel, manually extracted from 86 H&E-stained FFPE slides from the NCT Biobank (Heidelberg) and the UMM pathology archive (Mannheim). It is released in both Macenko-normalised and non-normalised (NONORM) form ^21,22^. Tiles were split 85% / 15% into training and internal-validation sets, stratified by class (random seed 42). CRC-VAL-HE-7K (7,180 tiles, 50 patients with no overlap with the training set, same tissue bank) and the Kather-2016 collection^33^ served as held-out test sets. The Kather-2016 classes were mapped onto the nine-class scheme (tumour, stroma, lymphocytes, debris, mucosa, adipose and empty mapped to TUM, STR, LYM, DEB, NORM, ADI and BACK). The 625 “complex stroma” tiles, which contain single tumour cells within stroma and have no clean counterpart, were excluded, leaving 4,375 tiles. Kather-2016 contains no mucus or muscle tiles, and its 150 px tiles were resized to 224 px for evaluation, which changes the effective magnification and may contribute to the lower accuracy on this set. All datasets are licensed CC-BY-4.0.

### Model formulation

A section imaged at *m* µm per pixel is first rescaled by the factor *m*/0.5 so that one pixel spans

0.5 µm, the training resolution. The rescaled image is partitioned into non-overlapping tiles *t*_*rc*_ of 224 × 224 pixels (112 µm) indexed by grid row *r* and column *c*; the last row and column are shifted inwards so that the image border is covered. For a pixel *p* with red, green and blue intensities *I*_*p,k*_, the saturation is

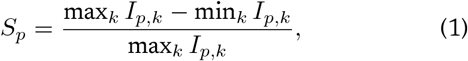

and a tile is treated as tissue, *τ* (*t*) = 1, unless more than 75% of its pixels have *S*_*p*_ *<* 0.10 (glass), in which case *τ* (*t*) = 0 and the tile is not scored.

Let *C* be the nine tissue classes (ADI, BACK, DEB, LYM, MUC, MUS, NORM, STR, TUM), *C* = |*C*| = 9. Each ensemble member *k* ∈ {1, 2, 3} is a convolutional network *g*_*k*_ whose ImageNet classification head was replaced by a linear layer with *C* outputs, giving class probabilities

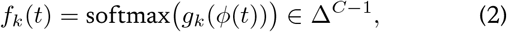

where *ϕ* standardises each colour channel with the ImageNet mean and standard deviation. The members are ConvNeXt-Tiny trained on Macenko-normalised tiles (*k* = 1), ConvNeXt-Tiny trained on NONORM tiles (*k* = 2) and EfficientNetV2-S trained on NONORM tiles (*k* = 3). At inference each tissue tile is first stain-normalised by the Macenko operator *N* (below), and the released model is the un-weighted average of member probabilities,

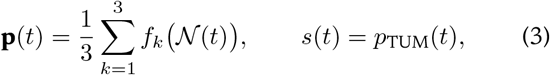

where *s*(*t*) ∈ [0, 1] is the tumour score of the tile. Averaging probabilities rather than logits keeps the combination insensitive to differences in the members’ logit scales. Supplementary Fig. S1 shows every step of this computation on a region of Cancer_P1.

### Model training

Members were initialised from ImageNet-pretrained weights distributed with the `timm` library ^56^ (`conv next_tiny.fb_in22k_ft_in1k` and `tf_efficientnetv 2_s.in21k_ft_in1k`; Apache-2.0) and fine-tuned end-to-end with PyTorch ^57^. For a mini-batch of tiles *x*_*i*_ with labels *y*_*i*_, the objective was class-weighted cross-entropy with label smoothing ^58^, as implemented in PyTorch,

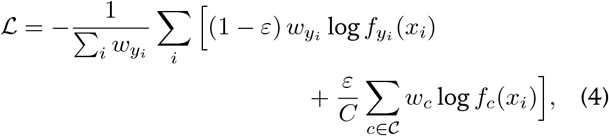

with smoothing *ε* = 0.1 and inverse-frequency class weights

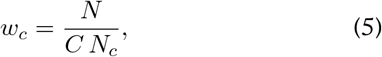

where *N*_*c*_ is the number of training tiles of class *c* and *N* the total. Parameters were optimised with AdamW ^59^ (weight decay 0.05) under a one-cycle schedule^60^ rising to a peak learning rate of 3 × 10^−4^ over the first 25% of steps and then annealing, for 8 epochs at batch size 128, with gradient-norm clipping at 1.0 and bfloat16 mixed precision. The checkpoint with the highest internal-validation balanced accuracy was retained. Augmentation targeted stain and orientation invariance: random resized crop (scale 0.7–1.0), horizontal and vertical flips, random 90^°^ rotations, colour jitter (brightness, contrast and saturation 0.25; hue 0.08) and random erasing (*p* = 0.25).

### Stain normalisation

We implemented the Macenko method ^26^. For each pixel, optical density is computed per channel as

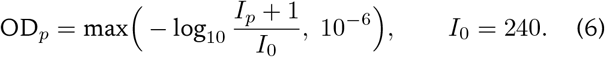

Pixels with optical density of at least *β* = 0.15 in all three channels are retained (at most 20,000, sampled at random). With **e**_1_, **e**_2_ the eigenvectors of their 3 × 3 covariance matrix with the largest and second-largest eigenvalues, each retained pixel is projected onto the plane they span and assigned the angle

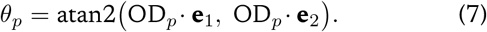

The directions at the 1st and 99th percentiles of *θ*, **u**(*θ*) = **e**_2_ cos *θ* + **e**_1_ sin *θ*, are taken as the two stain vectors. After unit normalisation they form the stain matrix **M** = [**h e**], with haematoxylin **h** the vector with the larger red-channel component. Stain concentrations are obtained by least squares, **c**_*p*_ = **M**^+^OD_*p*_, where **M**^+^ is the Moore–Penrose pseudo-inverse. Each stain’s concentrations are rescaled so that their 99th percentile matches a fixed reference **c**^ref^ = (1.9705, 1.0308), and the tile is reconstructed with the reference stain matrix

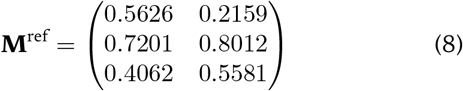

where 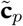 are the rescaled concentrations. Clamping the reconstructed optical density prevents numerical overflow on noisy tiles. Tiles for which the stain matrix cannot be estimated (fewer than 50 retained pixels, or a non-positive concentration percentile) are passed through unchanged. Normalisation adds about 10 ms per tile on a CPU.

### Tumour mask and region export

The tile-level tumour mask at threshold *ϑ* is

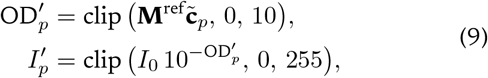

which is smoothed on the tile grid by a morphological closing followed by an opening with a 3 × 3 structuring element *K*,

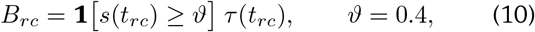

*B*^′^ is painted back onto the pixel grid, and its connected regions are traced as polygons with inner holes (OpenCV `findContours`). Regions smaller than *A*_min_ = 5 × 10^4^ px^2^ (12,500 µm^2^, about one tile) and holes smaller than *A*_min_/10 are discarded. Vertex coordinates are divided by the rescaling factor, so the GeoJSON polygons align with the image supplied by the user. The reported tumour area of a section is the fraction of tissue tiles above threshold before smoothing,

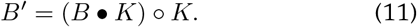

Unless stated otherwise, results for the released model use *ϑ* = 0.4. The comparison across inference variants (Fig. 1d; Supplementary Table S2) used *ϑ* = 0.5. Algorithm 1 summarises the procedure.

#### Algorithm 1.

**Tumour-region mapping of one section**

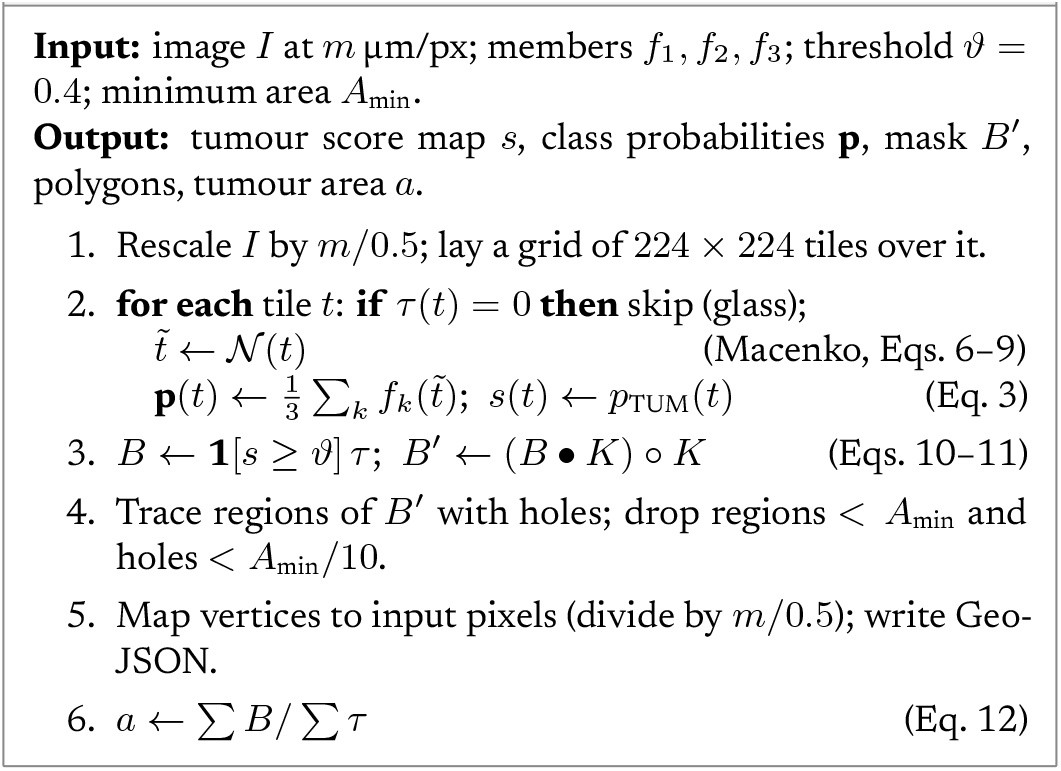

### Evaluation sections

The five evaluation sections are 10x Visium HD FFPE colorectal sections from Oliveira et al. ^6,34^: primary CRC from patients P1, P2 and P5 and normal-adjacent tissue from patients P3 and P5. H&E images were rescaled from their native resolution (0.274 µm per pixel) to 0.5 µm per pixel and cropped to the capture area plus a 150 µm border, using the bin2cell geometry^35^. Visium HD sections of lung adenocarcinoma (`Visium_HD_Human_Lung_Cancer_HD_Only_Experiment1`) and prostate adenocarcinoma (`Visium_HD_Human_Prostate_Cancer_FFPE`) from the 10x Genomics public dataset collection served as out-of-organ negative controls. None of these sections contributed to training or model selection.

### Transcriptomic reference

For Cancer_P1, cells had been reconstructed in a companion analysis by segmenting nuclei on the H&E image and assigning 2 µm bins to them with bin2cell ^35^. Each cell was then labelled by a multinomial logistic-regression classifier (L2 penalty, *C* = 0.1). The classifier was trained on the matched Chromium Flex single-cell reference of the same study ^6^, using 38 level-2 cell types, 1,246 marker genes and 79,573 training cells, with each training cell down-sampled to the sequencing depth of the spatial cells. Cells assigned to any of the tumour level-2 types formed the tumour lineage. When patient P1 was held out of training, the classifier assigned the correct broad lineage to 87.8% of P1 reference cells (level-2 accuracy 55.8%). Cell centroids were mapped into image coordinates using the recorded bin2cell scale factor, which equals the source resolution divided by 0.5 µm; all 202,182 cells were matched and fell within the scored grid. Tiles with at least 20 cells were scored, and a tile was called molecularly positive when at least half its cells were tumour-lineage.

### Statistics

For the transcriptomic comparison, each Cancer_P1 tile *t* containing *n*_*t*_ ≥ 20 cells was assigned a molecular tumour fraction 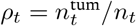 and a binary reference label *y*_*t*_ = **1**[*ρ*_*t*_ ≥ 0.5]. With *P* and *Q* the sets of positive and negative tiles, discrimination is summarised by the area under the ROC curve in its Mann–Whitney form,

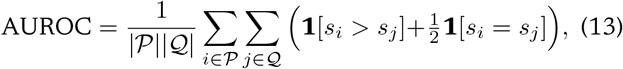

and, at threshold *ϑ*, by precision TP/(TP + FP), sensitivity TP/(TP + FN) and specificity TN/(TN + FP), where a tile is predicted positive when *s*_*t*_ ≥ *ϑ*. Rank agreement between *s*_*t*_ and *ρ*_*t*_ is Spearman’s *ρ*. Section-level separation between the carcinoma set *T* and the normal-adjacent set *R* is

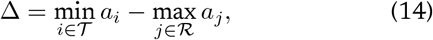

so that Δ *>* 0 means the two groups do not overlap.

Neighbouring tiles are spatially correlated, so tile-level resampling would understate uncertainty. We therefore used a spatial block bootstrap: the tile grid was divided into contiguous 8 × 8-tile blocks (1.8 mm across), the *B* = 57 non-empty blocks were resampled with replacement, all metrics were recomputed, and 95% intervals were taken as the 2.5th and 97.5th percentiles of 2,000 replicates. For comparison, a naive tile-level bootstrap gave a sensitivity interval of 0.902– 0.938 at threshold 0.4, against 0.845–0.966 for the block bootstrap. Confidence intervals for the held-out tile benchmarks, which pool tiles from many patients, use a tile-level bootstrap (1,000 resamples). The per-section results come from single slides and are reported descriptively, without inferential tests.

### Software and hardware

Python 3.11, PyTorch 2.11 (CUDA 12.8), timm 1.0.29, scikit-learn 1.9, OpenCV 5.0 and NumPy 2.4, on a single NVIDIA GeForce RTX 5080 Laptop GPU (16 GB). An auxiliary U-Net^61^ with a ResNet-34 encoder ^62^, trained on EBHI-SEG^63^ (validation IoU 0.866), is distributed with the package for boundary refinement but was not used for any result reported here.

## DATA AVAILABILITY

All training and test data are public: NCT-CRC-HE-100K, NCT-CRC-HE-100K-NONORM and CRC-VAL-HE-7K (doi:10.5281/zenodo.1214456); Kather-2016 (doi:10.5281/zenodo.53169); the Visium HD colorectal sections (doi:10.5281/zenodo.15042463); and the 10x Genomics lung and prostate Visium HD datasets (https://www.10xgenomics.com/datasets). Per-section outputs, summary tables and the per-tile agreement data underlying Figs. 2–4 are available from the corresponding author upon reasonable request.

## CODE AVAILABILITY

The OmiCoreTumorDetector package, including inference, training and evaluation code: https://github.com/OmiCore-Japan/omicoretumordetector. Package: https://pypi.org/project/omicoretumordetector/ (version 0.1.0). Model weights, with SHA-256 checksums: release v0.1.0 of the repository. Licence: Apache-2.0. The weights are derivative works of CC-BY-4.0 datasets. Other scripts used in this study, including those that generate the figures, are assets of OmiCore Inc. and are available from the corresponding author upon reasonable request.

## ETHICS STATEMENT

This study analysed only publicly available, de-identified datasets, downloaded from their public repositories and used under their licences. The data were collected by their original authors under those studies’ own ethics approvals; for example, the NCT-CRC-HE-100K data were collected under approval of Ethics Board II at University Medical Center Mannheim (2017-806R-MA) ^22^. The authors collected no new human samples or data, had no access to identifying information, and did not submit an application to an ethics committee, as analysis of published, de-identified data does not require one.

## AUTHOR CONTRIBUTIONS

N.S.: conceptualisation, methodology, software, validation, formal analysis, writing – original draft. A.F.: supervision, writing – review and editing.

## USE OF AI-ASSISTED TOOLS

The manuscript was drafted with open-source large language models, run locally and augmented by retrieval (retrieval-augmented generation, RAG) over research literature in the field, and was then extensively edited by N.S. and A.F. Software development, data analysis, figure generation and manuscript preparation were also assisted by a largelanguage-model coding assistant (Claude Code, Anthropic). The manuscript was typeset with LATEX, and all figures were generated with Python. The authors reviewed, verified and take full responsibility for all code, results and text.

## ACKNOWLEDGEMENTS

We thank J. N. Kather and colleagues for releasing the NCT-CRC-HE-100K, CRC-VAL-HE-7K and Kather-2016 datasets, the authors of Oliveira et al. and 10x Genomics for releasing the Visium HD colorectal data, and the authors of EBHI-SEG.

## SUPPLEMENTARY INFORMATION

**Table S1.** Held-out tile benchmarks for the individual ensemble members. Tumour AUROC is tumour-versus-rest, with 95% bootstrap intervals. Kather-2016 is balanced across its classes, so accuracy equals balanced accuracy.

| Model | CRC-VAL-HE-7K (7,180 tiles) |  |  | Kather-2016 (4,375 tiles) |  |  |
| --- | --- | --- | --- | --- | --- | --- |
|  | Acc. | Bal. acc. | Tumour AUROC | Acc. | Bal. acc. | Tumour AUROC |
| ConvNeXt (normalised) | 0.970 | 0.955 | 0.999 (0.998–1.000) | 0.569 | 0.569 | 0.836 (0.813–0.859) |
| ConvNeXt (NONORM) | 0.780 | 0.756 | 0.987 (0.985–0.990) | 0.763 | 0.763 | 0.992 (0.986–0.997) |
| EfficientNetV2 (NONORM) | 0.821 | 0.801 | 0.994 (0.992–0.997) | 0.772 | 0.772 | 0.994 (0.989–0.998) |

**Table S2.** All inference variants at section level and against the transcriptomic reference. Section values are mean tumour area (%) at threshold 0.5. Separation is the smallest carcinoma-section area minus the largest normal-adjacent area, in percentage points. Out-of-organ: mean of the lung and prostate sections. AUROC: agreement with transcriptomic identities on Cancer_P1. TTA, test-time augmentation over four flips.

| Variant | Carcinoma | Normal adj. | Out-of-organ | Separation | AUROC |
| --- | --- | --- | --- | --- | --- |
| Ensemble + stain norm (released) | 34.1 | 0.57 | 3.3 | +24.6 | 0.985 |
| ConvNeXt + stain norm | 21.9 | 0.17 | 1.0 | +18.1 | 0.955 |
| Ensemble | 31.0 | 1.14 | 3.7 | +16.4 | 0.983 |
| ConvNeXt (NONORM tiles) | 35.1 | 3.06 | 7.3 | +14.2 | 0.979 |
| ConvNeXt + TTA | 22.2 | 2.54 | 4.2 | +12.7 | 0.960 |
| ConvNeXt (normalised tiles) | 22.2 | 2.92 | 4.9 | +11.5 | 0.955 |
| EfficientNetV2 (NONORM tiles) | 30.0 | 1.27 | 6.1 | +10.7 | 0.971 |

**Table S3.** Sensitivity of the molecular comparison to the ground-truth definition. The minimum number of cells per tile and the tumour-cell fraction required to call a tile molecularly positive were varied. AUROC of the released model’s tumour probability.

| Min. cells | Tumour fraction | Tiles | Positive tiles | AUROC |
| --- | --- | --- | --- | --- |
| ≥20 | ≥0.3 | 2,307 | 938 | 0.9774 |
| ≥20 | ≥0.5 | 2,307 | 859 | 0.9855 |
| ≥20 | ≥0.7 | 2,307 | 758 | 0.9823 |
| ≥40 | ≥0.3 | 1,936 | 846 | 0.9846 |
| ≥40 | ≥0.5 | 1,936 | 778 | 0.9900 |
| ≥40 | ≥0.7 | 1,936 | 692 | 0.9807 |

**Figure S1.**
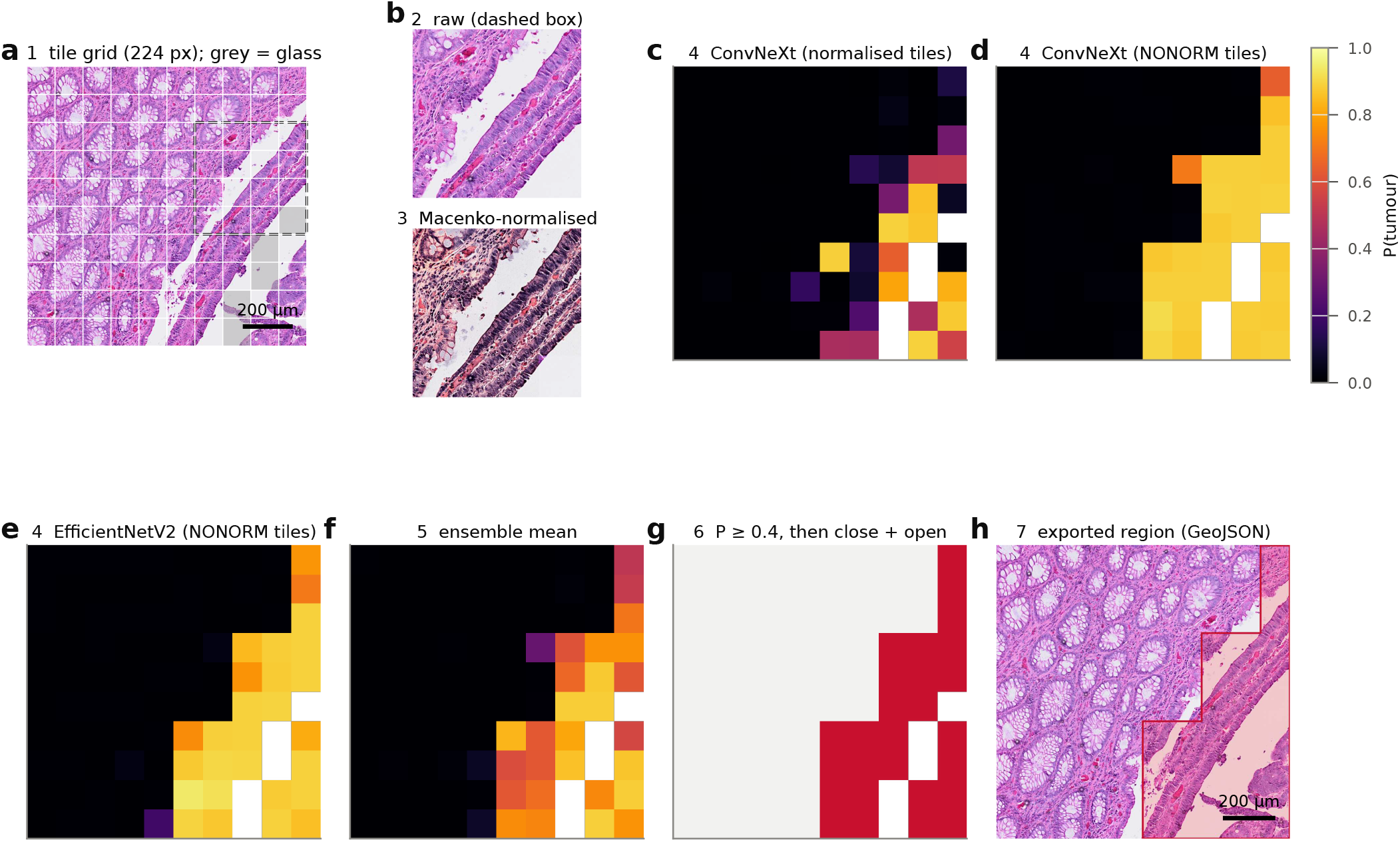
Step-by-step inference on one region. A 10 × 10-tile region (2,240 × 2,240 px, 1.12 mm across) of Cancer_P1 that straddles the tumour boundary, processed exactly as by the released model (Algorithm 1). **a**, H&E region at 0.5 µm per pixel with the 224-px tile grid. Grey tiles were classified as glass (more than 75% of pixels with saturation below 0.10; Eq. 1) and were not scored. The dashed box marks the 4 × 4-tile block shown in **b. b**, The block before (top) and after (bottom) Macenko normalisation to the reference stain profile (Eqs. 6–9); normalisation is applied to each tile independently. **c– e**, Tumour probability from each ensemble member applied to the normalised tiles: ConvNeXt-Tiny trained on normalised tiles (**c**), ConvNeXt-Tiny trained on non-normalised tiles (**d**) and EfficientNetV2-S trained on non-normalised tiles (**e**). The members largely agree but differ in confidence near the boundary. **f**, Ensemble mean of **c–e**, which is the released tumour score (Eq. 3). It reproduces the whole-section ensemble map to within 0.011, a difference due to reduced-precision (bfloat16) arithmetic in the released pipeline. **g**, Tumour mask after thresholding at 0.4 and a 3 × 3 morphological closing and opening on the whole-section grid (Eqs. 10–11); dark red, tumour; light grey, non-tumour tissue; white, glass. Smoothing changed no tiles in this region. **h**, The exported GeoJSON region (red outline, light tint) over the H&E region. The colour scale at right applies to **c–f**; white tiles in **c–f** are glass (not scored). Scale bars, 200 µm. Images are embedded at 1,000–1,477 ppi.

